# Integrated Regulation of PKA by Fast and Slow Neurotransmission in the Nucleus Accumbens Controls Plasticity and Stress Responses

**DOI:** 10.1101/2021.09.29.462408

**Authors:** Rachel Thomas, Adan Hernandez, David R. Benavides, Wei Li, Chunfeng Tan, Florian Plattner, Ayanabha Chakraborti, Lucas Pozzo-Miller, Susan S. Taylor, James A. Bibb

## Abstract

Cortical glutamate and midbrain dopamine neurotransmission converge to mediate striatum-dependent behaviors, while maladaptations in striatal circuitry contribute to mental disorders. Here we uncover a molecular mechanism by which glutamatergic and dopaminergic signaling integrate to regulate cAMP-dependent protein kinase (PKA) via phosphorylation of the PKA regulatory subunit, RIIβ. We find that glutamate-dependent reduction in Cdk5-dependent RIIβ phosphorylation alters the PKA holoenzyme auto-inhibitory state to increase PKA signaling in response to dopamine. Disruption of RIIβ phosphorylation by Cdk5, consequently, enhances cortico-ventral striatal synaptic plasticity. Acute and chronic stress in rats inversely modulate RIIβ phosphorylation and ventral striatal infusion of a small interfering peptide that selectively targets RIIβ regulation by Cdk5 improves behavioral response to stress. This new signaling mechanism integrating ventral striatal glutamate and dopamine neurotransmission is likely important to brain function, may contribute to neuropsychiatric conditions, and serves as a possible target for the development of novel therapeutics for stress-related disorders.

## INTRODUCTION

A fundamental function of brain circuitry is to impart emotional salience upon sensory input so that environmental experiences may be acted upon with appropriate behavioral responses. This requires the integration of fast ligand-gated ionotropic synaptic signals with those mediated by G-protein coupled neurotransmitter receptors. Subserving these synaptic receptors, Ca^2+^ and cAMP are two principle second messengers that invoke overlapping intracellular signaling networks which compute the appropriate responses in the form of altered excitability and synaptic strength upon which learned behavioral responses are based. Understanding of the molecular mechanisms by which this is achieved is incomplete.

The nucleus accumbens (NAc, ventral striatum) is a central input structure within the basal ganglia and mediates reinforced learning and behavioral response processing. The medium spiny neurons (MSNs) in the NAc receive glutamatergic input from the medial prefrontal cortex, hippocampus, and other regions as well as dopaminergic input from the ventral tegmental area (VTA) (Salgado and Kaplitt, 2015, Xu et al., 2020). Coincidental detection of these two inputs is critical to reward, aversion, motor learning, and other aspects of sensory and emotional integration and behavioral flexibility (Uddin, 2021, Miyazaki et al., 2013). The contributions of glutamate and dopamine to these critical brain functions are mediated by the interplay of key signaling pathways, such as the PKA cascade (Yagishita et al., 2014, Greengard et al., 1999, Yapo et al., 2018).

The tetrameric PKA holoenzyme consists of two catalytic domains (PKA_cat_: Cα, Cβ, or Cγ isoforms) bound to a homodimer of regulatory subunits (RIα, RIβ, RIIα, or RIIβ isoforms). When RI subunits are incorporated, it is classified as Type I PKA; when bound to RII subunits as Type II PKA. R subunits possess an N-terminal dimerization domain (D/D), responsible for both dimerization and subcellular localization via binding to members of the A-kinase anchoring protein (AKAP) family of scaffolding molecules (Omar and Scott, 2020). Each R subunit monomer contains two C-terminal cyclic nucleotide-binding (CNB) domains. Upon cAMP binding, PKA is activated by unleashing the catalytic activity from R dimer inhibition (Taylor et al., 2013). Between the D/D and CNB domains is a flexible linker region containing the inhibitor sequence (IS) responsible for occupying the C subunit active site in the inhibited state under basal conditions. The linker imparts an important structural distinction between Type I and Type II PKA, with functional implications for the holoenzyme. While the IS in Type I R subunits is a pseudosubstrate for PKA, the Type II IS acts as a true substrate for the C subunits at Ser95 on RIIα and Ser114 on RIIβ (Zhang et al., 2012). Phosphorylation at the PKA site on RII subunits has been suggested to slow the reassociation rate of the R and C subunits (Rangel-Aldao and Rosen, 1976, Isensee et al., 2018).

PKA activation engages many downstream effectors and signaling cascades, and many pathways impinge upon cAMP/PKA signaling. The cAMP/PKA cascade may be activated by either ligand-gated ionotropic or G_s_ protein receptor-coupled metabotropic neurotransmission (Greengard et al., 1999, Sassone-Corsi, 2012). Previously we showed that constitutive phosphorylation of DARPP-32 (Bibb et al., 1999, Nishi et al., 2000) by Cdk5 serves as an important mechanism by which glutamate and dopamine interact to regulate PKA signaling. Also, we reported that PKA-dependent activation of cAMP-specific phosphodiesterases is modulated though Cdk5-dependent phosphorylation of PDE4 family members (Plattner et al., 2015). However, *direct* regulation of the PKA holoenzyme has not previously been shown to fall under the coordinated control of excitatory and metabotropic neurotransmission. We hypothesized that control of PKA activation/inhibition could serve as a critical point of integration for glutamate and dopamine inputs. Here we report a novel mechanism by which these two modes of neurotransmission are integrated to mediate ventral striatal circuitry function in plasticity and stress-response behavior.

## RESULTS

### The PKA regulatory subunit RIIβ is a substrate for Cdk5

Messenger RNA transcripts encoding all four isoforms of the PKA regulatory subunit are present in brain, with expression of the RIIβ isoform enriched in CNS (Cadd and McKnight, 1989, Ventra et al., 1996, Ilouz et al., 2012). Examination of protein expression (**Figure 1A**) indicates that the RIIβ subunit is most selectively expressed in brain compared to the other regulatory isoforms. Moreover, its brain region tissue distribution mimics the expression pattern of Cdk5. For both RIIβ and Cdk5, highest levels occur in limbic regions including prefrontal cortex, dorsal and ventral striatum, and hippocampus. Immunostains of ventral striatum showed RIIβ in the soma and neuropil of MSNs where it colocalized with its scaffolding protein, AKAP150 (Carr et al., 1992, Murphy et al., 2014) (**Figure 1B**). RIIβ also co-localized with Cdk5 in dissociated rat striatal neurons grown in culture (**Figure 1C**).

**Figure 1.**
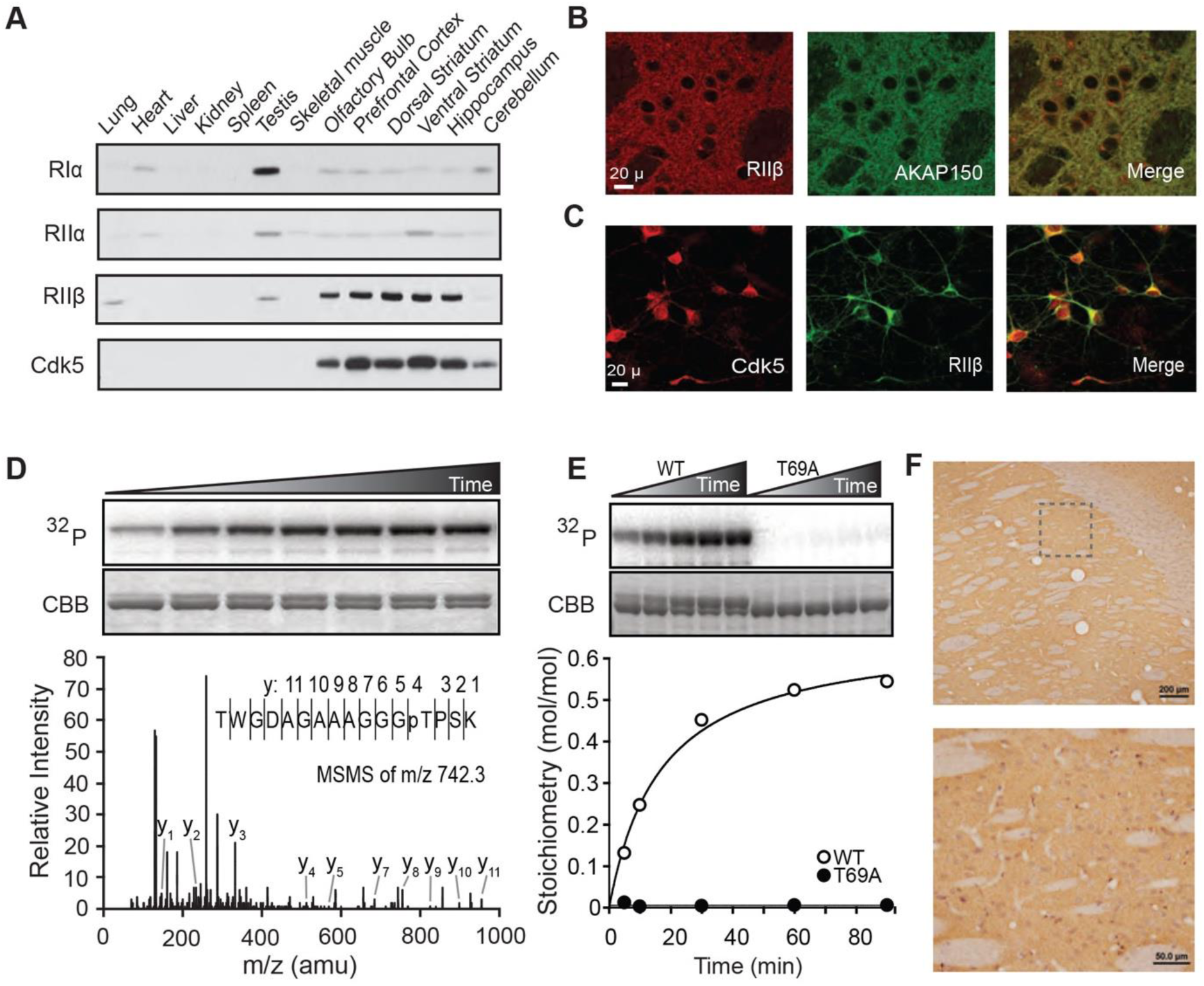
PKA regulatory subunit RIIβ phosphorylation at Thr69 by Cdk5. (A) Tissue distribution of R subunits and Cdk5 in rat peripheral and CNS tissues. (B) Co-immunostain of RIIβ and AKAP150 in rat ventral striatum. (C) Co-localization of Cdk5 and RIIβ in dissociated rat striatal neurons. (D) Time-course *in vitro* phosphorylation of RIIβ by Cdk5 (autoradiogram, ^32^P, top; Coomassie stain CBB, middle) with accompanying ESI-Qq TOF MS/MS spectrum (bottom) of the tryptic peptide shown encompassing phospho-Thr69 (pT, ion peak y4) positively identifying the site of RIIβ phosphorylation by Cdk5. (E) *In vitro* phosphorylation of WT vs. T69A RIIβ by Cdk5 with stoichiometry. (F) Immunostain of phospho-Thr69 RIIβ (brown) in neuropil with nuclear counterstain (purple) in rat striatum. Box insert (top) indicates region shown at higher magnification (bottom). See also Figure S1.

The amino acid sequence of RIIβ includes a proline-directed Cdk5 consensus motif (S/TPXK/H/R) within the N-terminal linker region predicting Thr69, a site unique to this particular R subunit, as a potential Cdk5 phosphorylation site. Indeed, Cdk5 phosphorylated pure recombinant RIIβ *in vitro*, achieving a maximal stoichiometry of 0.6 mol PO_4_/mol of substrate under saturating conditions (**Figures 1D and 1E**). Mass spectrometry positively identified Thr69 as the site of phosphorylation, and the T69A site-directed mutant form of RIIβ was not phosphorylated by Cdk5 at all. To determine if this phosphorylation occurred in brain, a phosphorylation state-specific antibody for phospho-Thr69 RIIβ was derived (**Figure S1A-S1C**). Phospho-Thr69 RIIβ was detected within MSNs and neuropil throughout striatum by immunostaining. It was not detected in oligodendrocytes or adjacent white matter (**Figure 1F**). Together these data demonstrate that the RIIβ regulatory subunit of PKA is phosphorylated at Thr69 by Cdk5, and that this phosphorylation event occurs *in vivo* within striatal neurons.

### Phosphorylation of RIIβ by Cdk5 affects PKA holoenzyme autophosphorylation kinetics

Both genetic deletion and pharmacologic inhibition of Cdk5 increases phosphorylation of PKA substrates in striatum (Plattner et al., 2015). Thus, we hypothesized that Cdk5-mediated phosphorylation of Thr69 RIIβ could have an inhibitory effect on PKA_cat_ activity. However, *in vitro* analysis indicated that Thr69 phosphorylation neither altered the inhibition of PKA catalytic activity by RIIβ (**Figure S1D**), nor cAMP-dependent activation of PKA in the RIIβ/PKA holoenzyme complex (**Figure S1E**). As an alternative, we considered that Cdk5-dependent phosphorylation of Thr69 RIIβ could alter structure/function aspects of the holoenzyme inhibitory state. As mentioned above, the RIIβ dimerization domain forms a four-helix bundle that docks with AKAPs through hydrophobic interactions (Esseltine and Scott, 2013, Kinderman et al., 2006) (**Figure 2A**). The Cdk5 phosphorylation site on RIIβ is encompassed within the flexible linker region that occurs between the dimerization domain and the cAMP binding domains. This site is proximal to the inhibitor sequence in Type II R subunits, which contains the PKA phosphorylation site Ser114 (**Figure 2B**). Therefore, we assessed the effect of phospho-Thr69 on PKA-dependent phosphorylation of Ser114 RIIβ (**Figure 2C**). Interestingly, T69D phospho-mimetic mutation attenuated the efficiency of Ser114 phosphorylation by PKA in comparison to WT RIIβ, significantly reducing the maximum velocity of the reaction under linear conditions. Thus, the phosphorylation state of RIIβ at Thr69 governs PKA “autophosphorylation” at Ser114.

**Figure 2.**
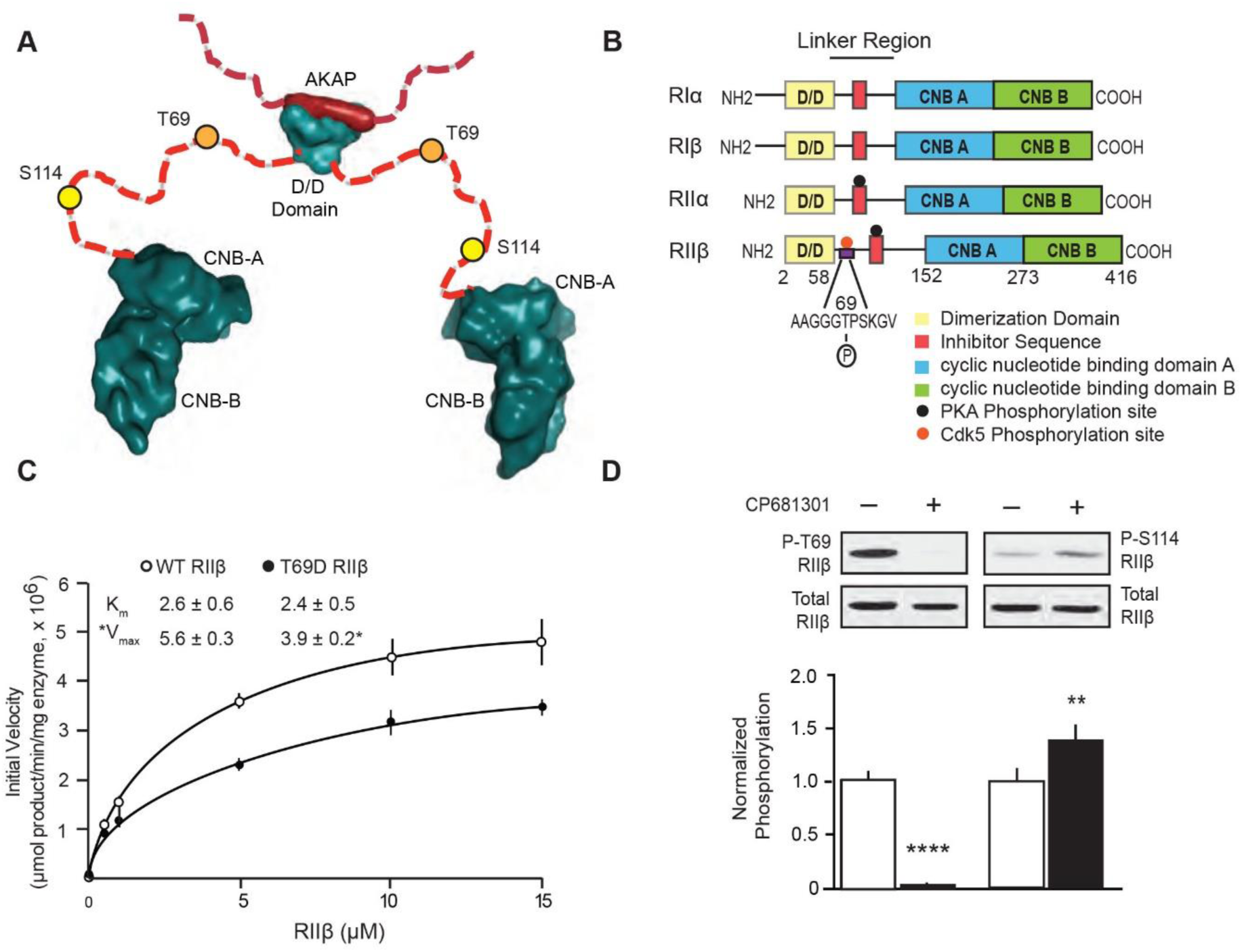
The phosphorylation state of Thr69 governs PKA-dependent phosphorylation of Ser114 RIIβ. (A) Structural model of RIIβ dimer complexed with AKAP showing the flexible linker region with Cdk5 and PKA sites denoted (Diller et al., 2001, Kinderman et al., 2006). (B) Regulatory subunit isoforms of PKA with positions of structural/functional domains and phosphorylation sites indicated. (C) Kinetic analysis of *in vitro* auto-phosphorylation of Ser114 WT vs. T69D RIIβ by PKA_cat_ with K_m_ and V_*max*_ values. Data represent means ± S.E.M., \**p*<0.05, unpaired *t*-test, n=3. (D) Effects of striatal slice treatment with the Cdk5 inhibitor, CP681301 on phosphorylation of RIIβ at Thr69 (P-T69) by Cdk5 and Ser114 (P-S114) by PKA. Immunoblots of lysates from striatal slices treated with CP681301 (50 µM, 1 h) are shown with quantitation. All data are normalized means ± S.E.M., \*\*\*\**p*<0.0001 for P-T69, \*\**p*<0.01 for P-S114 RIIβ, unpaired *t*-test, n=8-9. See also Figure S2.

To determine if this functional relationship between Thr69 and Ser114 RIIβ phosphorylation occurs *ex vivo*, acute striatal brain slices were treated with the Cdk5 inhibitor, CP681301 (**Figure 2D**). Cdk5 inhibition attenuated the phosphorylation of phospho-Thr69, with corresponding increase in PKA-dependent phosphorylation of Ser114. Together, these data reveal a novel intramolecular mechanism by which RIIβ phosphorylation is reciprocally regulated at two proximal sites *in vitro* and *ex vivo*.

### Mechanistic function and regulation of RIIβ/PKA in ventral striatum

PKA regulation via RII subunit autophosphorylation has not previously been examined as a neuronal signaling mechanism. This may be because phosphorylation of RII by PKA characterizes the inhibited state of type II PKA holoenzymes (Zhang et al., 2012). However, early studies suggested that phosphorylation at the PKA site could slow RIIβ reassociation with PKA_cat_ following holoenzyme dissociation (Rangel-Aldao and Rosen, 1976). Consistent with this mechanistic function, phospho-mimetic S114D mutation of RIIβ reduced the PKA inhibition potency over 4-fold (IC_50_ = 1.5 ± 0.3 nM, WT; 6.1 ± 1.1, S114D) (**Figure 3A**). Thus, the level of PKA-dependent phosphorylation of Ser114, which occurs with formation of the PKA/RIIβ holoenzyme inhibitory state is controlled by the phosphorylation state of Thr69 RIIβ and may determine PKA inhibition potency by RIIβ. Therefore, neuronal activity which regulates Thr69 levels could modulate the efficacy of activators of G protein-coupled receptors that invoke the cAMP/PKA signaling cascade.

**Figure 3.**
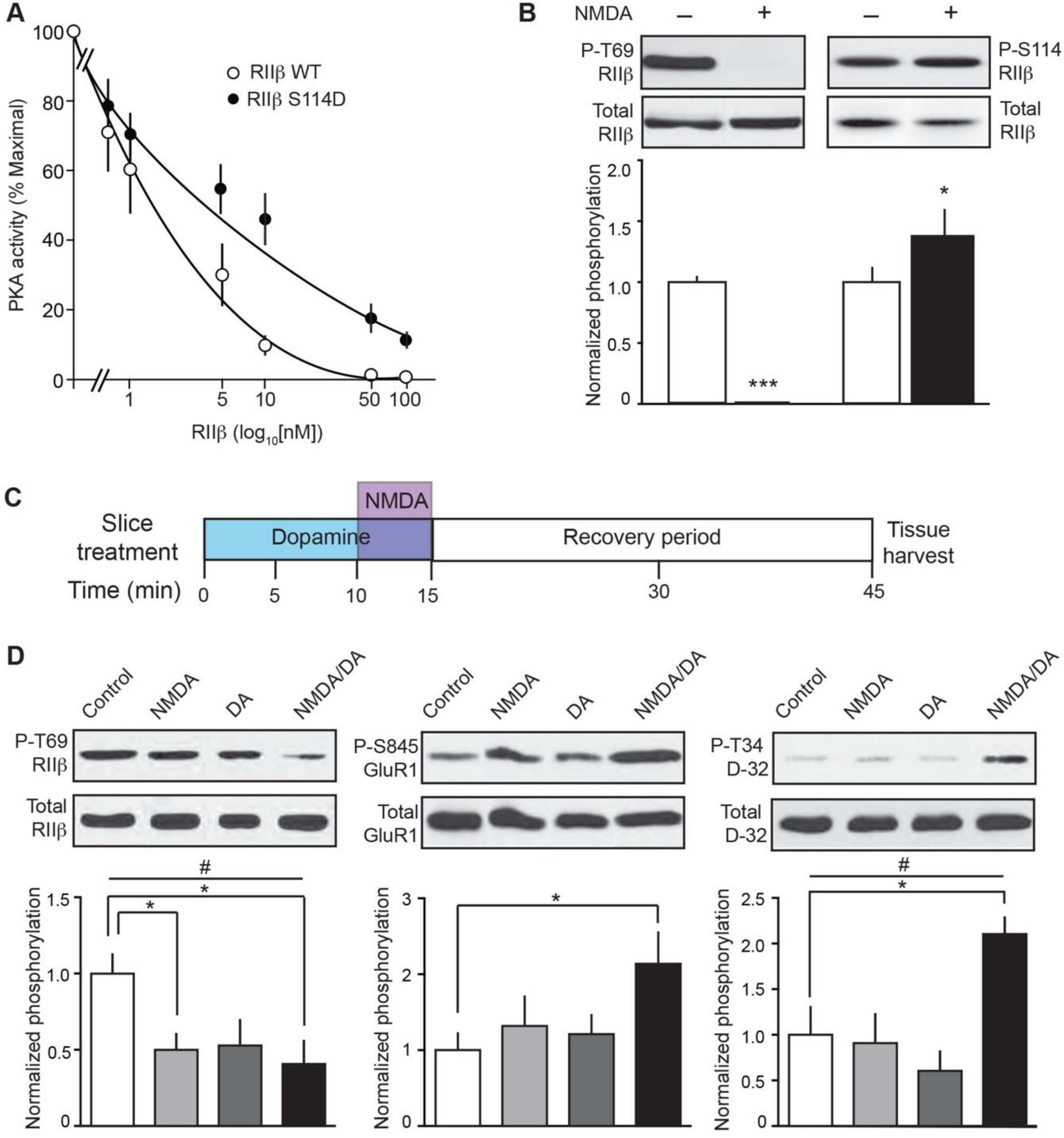
Reciprocal regulation of proximal RIIβ phosphorylation sites allows integration of glutamatergic and dopaminergic neurotransmission. (A) PKA inhibition curves for WT vs. S114D RIIβ. Data represent means ± S.E.M., IC_50_ = 6.1 ± 1.1, S114D; 1.5 ± 0.3 nM, WT, \**p*<0.05, unpaired *t*-test, n=3. (B) Quantitative immunoblot analysis of lysates from striatal slices treated with NMDA (50 µM, 5 min) showing reciprocal regulation of the phospho-Thr69 and Ser114 RIIβ. Data represent means ± S.E.M., \*\*\**p*<0.001 for P-T69, \**p*<0.05 for P-S114, unpaired *t*-test, n=4. (C) Schematic of NMDA/dopamine slice pharmacology co-treatment paradigm. (D) Effects of striatal slice treatment with NMDA (25 µM, 5 min), dopamine (DA, 10 µM, 15 min), or both on Cdk5-dependent phosphorylation of Thr69 RIIβ (^#^*p*<0.05, one-way ANOVA; \**p*<0.05, multiple comparisons, control vs. NMDA and control vs. NMDA/DA, n=6), PKA-dependent phosphorylation of Ser845 GluA1 (\**p*<0.05, multiple comparisons, control vs. NMDA/DA, n=5-6) and Thr34 DARPP-32 (^##^*p*<0.005, one-way ANOVA; \**p*<0.05, multiple comparisons, control vs. NMDA/DA, n=6). Data represent means ± S.E.M.

To further assess this possibility, the regulation of phospho-Thr69 and Ser114 RIIβ was explored in striatal slices. Acute NMDA treatment (50 µM, 5 min) strongly reduced the phosphorylation state of Thr69 (**Figure 3B**). Concomitantly, PKA-dependent phosphorylation at Ser114 was increased, consistent with the reciprocal modulation of these sites detected *in vitro* and in response to Cdk5 inhibition in intact brain tissue. The effect of NMDA on phospho-Thr69 appeared to be mediated by the activation of the Ca^2+^-dependent serine/threonine protein phosphatase, PP2B (calcineurin), as the NMDA-induced reduction was blocked by the PP2B inhibitor cyclosporin A (**Figure S2A**). Inhibition of protein phosphatases PP1 and PP2A by 1 µM okadaic acid also attenuated the effect of NMDA. Moreover, the basal phosphorylation state of phospho-Thr69 was increased by PP2B inhibition with cyclosporin A, PP1 inhibition by 1 µM okadaic acid, or selective PP2A inhibition with 200 nM okadaic acid (**Figure S2B**). Together these data indicate that phospho-Thr69, and thereby phospho-Ser114, are regulated by activation of ionotropic NMDA class glutamate receptors. This is mediated, at least in part, through Ca^2+^-dependent protein phosphatases including PP2B and 2A. These phosphatases together with PP1 also determine the basal phosphorylation state of phospho-Thr69 RIIβ. The diversity in protein phosphatases dephosphorylating RIIβ may not be surprising as both RII subunits interact with the serine/threonine phosphatase calcineurin (PP2B), Cdk5 complexes with PP2A (Plattner et al., 2006), and AKAP signalosomes cluster protein phosphatases (Wong and Scott, 2004).

Phosphorylation at the Thr69 site modulates phosphorylation at Ser114, which in turn affects PKA inhibition by dissociated RIIβ. Therefore, NMDA-dependent modulation of Thr69 has the potential to impart effects upon PKA activation. To test this, ventral striatal slices were treated with NMDA to activate fast neurotransmission, dopamine to activate slow neurotransmission, or both. As both NMDA and dopamine receptor activation are essential to the induction of striatal LTP (Hernandez et al., 2016), these treatments were conducted under conditions similar to those used to induce striatal plasticity (**Figure 3C**). Specifically, dopamine (10 µM) was bath-applied for 15 min with NMDA (25 µM) added for the last 5 min in reduced Mg^2+^ buffer with a 30 min delay following treatments. Effects on phospho-Thr69 RIIβ compared with PKA activity, as indicated by changes in PKA-dependent phosphorylation states of two key effectors, Ser845 GluA1 and Thr34 DARPP-32 (**Figure 3D**). Under these conditions, NMDA again induced a decrease in phospho-Thr69 RIIβ, without significantly altering PKA activity. Dopamine alone also did not sustain PKA activity for 30 min. However, phospho-Thr69 reduction by combined dopamine and NMDA treatment corresponded to increased PKA activity 30 min after the treatments were removed (**Figure 3D**). These data suggest a compound mechanism in which relatively high levels of phospho-Thr69 occur under basal conditions. Robust excitatory glutamatergic neurotransmission reduces phospho-Thr69 via protein phosphatase activation, resulting in increased PKA-dependent phosphorylation of Ser114 RIIβ. In the context of dopamine receptor activation, this leads to sustained elevated PKA activity involving downstream effector signaling pathways known to mediate synaptic plasticity and striatal function.

### RIIβ phosphorylation controls ventral striatal plasticity

To directly assess the contribution of the RIIβ regulatory phosphorylation mechanism to striatal function, a small interfering peptide (siP) was developed which corresponded to 18 amino acid residues from the RIIβ linker region encompassing Thr69 and included a membrane-permeabilizing penetratin tag (**Figure 4A, top**). This Thr69 RIIβ siP (RIIβ siP) potently inhibited Cdk5-dependent phosphorylation of RIIβ (IC_50_ = 7.14 ± 2.28 µM) (**Figure S3A**). Treatment of ventral striatal slices with this peptide caused reduction in phospho-Thr69 RIIβ, but had no effect on Cdk5-dependent phosphorylation of Thr75 DARPP-32 (**Figure 4A, bottom**). RIIβ siP/dopamine co-treatment of ventral striatal slices resulted in a sustained increase in PKA activity, similar to the previously observed effects of NMDA/dopamine treatment (**Figure S3B**). Consequently, we established the means to selectively modulate phospho-Thr69 RIIβ levels in intact brain tissue and facilitate prolonged PKA activation.

**Figure 4.**
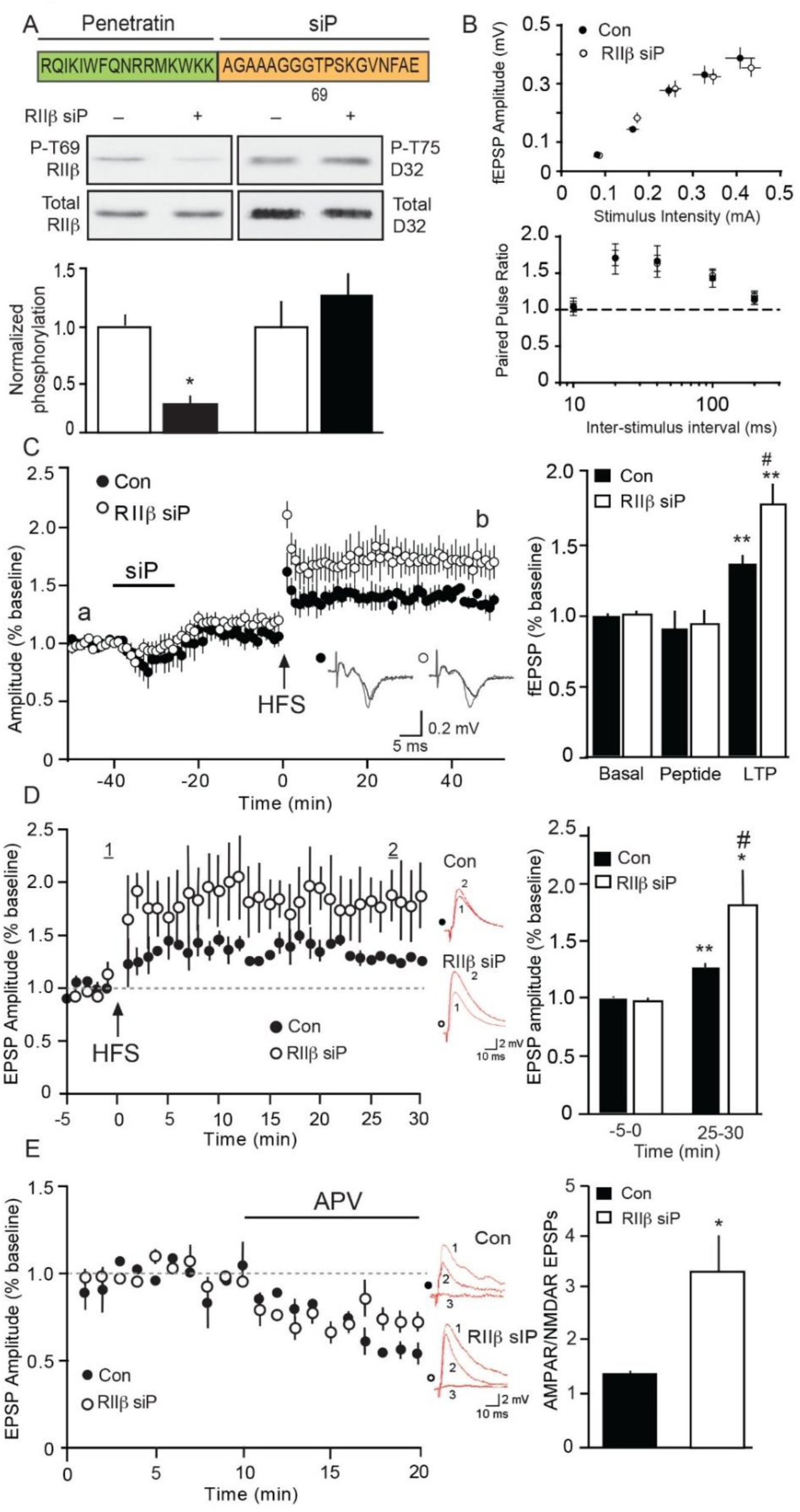
Selective interference of Thr69 RIIβ phosphorylation enhances ventral striatal plasticity. (A) Sequence of the RIIβ-targeting siP (top) and immunoblots of lysates from NAc slices treated with Th69 RIIβ siP (10 µM, 1 h). Data represent means ± S.E.M., \**p*<0.05, unpaired *t*-test, n=4. See also Figure S3. (B) Input/output (I/O) curves and paired pulse ratio (PPR) derived from cortico-ventral striatal fEPSP recordings of scrambled peptide control (Con)- and RIIβ siP-treated slices. (C) Effects of RIIβ siP on NAc LTP assessed by field recordings. Plot of fEPSP amplitudes with sample traces from a vs. b are shown (left) with summary plot (right) (^#^*p*<0.05 compared to scrambled peptide control, unpaired *t*-test; \*\**p*<0.01 compared to basal, Wilcoxon test, n=5). See also Figure S4. (D). Effect of RIIβ siP on LTP in D1R neurons. EPSP amplitude following HFS of cortico-ventral striatal circuitry (top). Inset shows representative traces of EPSPs in current-clamp mode before and 25 min post-LTP induction. Summary data (right) shows effect of RIIβ siP (10 µM, 1 h) vs control (scrambled siP). Data represent means ± S.E.M.; \*\**p* <0.01 compared to control baseline, \**p*<0.05, compared to RIIβ siP baseline, #*p*<0.05, compared to LTP in controls, Student’s *t* test, n = 4-6. (E) Effect of RIIβ siP on EPSP amplitude and response to selective ionotropic glutamate receptor antagonists in D1R neurons. Control and RIIβ siP-treated NAc slices underwent NMDAR antagonist APV (20 μM) treatments. Subsequent treatments with the AMPAR antagonist NBQX (10 μM) ablated EPSP responses (see inset). EPSP recordings (30 min after HFS, LTP induction) and APV effects are shown. Insets show traces for D1R neurons before (1) and after APV (2), and combined APV/NBQX (3) treatments. Summary data is presented (right) for AMPA/NMDA ratios. Data are means ± SEM, *p*\*<0.05, Student’s *t* test n = 4-6.

With this reagent in hand, we chose to assess the role of RIIβ phosphorylation in cortico-ventral striatal plasticity as one measure of physiological function. Cortical afferents within corpus callosum of sagittal brain slices were stimulated and field excitatory post-synaptic potentials (fEPSPs) were recorded in ventral striatum core (**Figure S4A**). Long-term potentiation (LTP) was induced by high frequency stimulation (HFS), and could be enhanced by 15 min bath application of the D1-type dopamine receptor agonist SKF81297 (**Figure S4B**), as previously observed for dorsolateral striatum (Hernandez et al., 2016). To assess the contribution of RIIβ phosphorylation to LTP, equilibrated slices were treated with either the RIIβ siP or a scrambled siP (control, 10 µM) for 15 min, followed by drug washout for 25 min prior to HFS (**Figures 4B and 4C**). The RIIβ siP had no effect on synaptic excitation (input-output, I/O) or paired pulse response ratios (PPR). RIIβ siP caused a transient reduction in basal fEPSP amplitude but significantly enhanced LTP compared to scrambled control, siP causing elevations in fEPSP amplitudes that were maintained through 50 min following HFS (0.31 ± 0.03 mV scrambled siP vs. 0.45 ± 0.06 mV for RIIβ siP). Interestingly, the enhancement of LTP by the RIIβ siP was partly occluded in slices pre-treated with SKF-81297 (2 µM) prior to HFS (**Figures S4C and S4D**).

To better understand this effect, whole cell patch-clamp recordings were made of D1R-expressing neurons (Ade et al., 2011). As observed in field recordings, the RIIβ siP induced a significant increase in EPSP amplitude and markedly elevated LTP levels in comparison to control treated neurons (**Figure 4D**). This effect was attributable to increased AMPAR-mediated function, as AMPA/NMDA ratios were increased by the RIIβ siP (**Figure 4E**), consistent with elevated phospho-Ser845 GluR1, which is linked to increased synaptic surface AMPAR trafficking (Henley and Wilkinson, 2013) Thus, direct siP targeting of Thr69 RIIβ phosphorylation enhances cortico-ventral striatal plasticity, likely by increasing PKA activity. These findings together with the slice signaling data (**see Figure 3**) are consistent with the modulation of RIIβ phosphorylation and PKA activity by NMDA and dopamine as an important mechanism mediating ventral striatal plasticity.

### RIIβ regulation is altered by stress and can be targeted to affect stress-related behaviors

Synaptic plasticity and cognition are affected by stress (Russo and Nestler, 2013, Dwivedi and Pandey, 2000, Aceto et al., 2020). While acute stress can improve cognitive performance, exposure to persistent stress causes cognitive impairment (Yuen et al., 2012, Bellani et al., 2006). To determine if stress could alter the signaling state of RIIβ phosphorylation, rats were exposed to a stressful environment either once (acute stress, one forced swim) or chronically (chronic unpredictable stress, CUS, two stressors daily for 14 days). Interestingly, one hour following exposure to acute stress, phospho-Thr69 RIIβ was significantly decreased (**Figure 5A**). In contrast, chronic exposure to stress caused elevation in the phosphorylation state of this site. Thus, environmental exposure to stressful conditions can alter the basal signaling state of RIIβ, raising the intriguing possibility that the regulation of RIIβ phosphorylation may contribute to behavioral responses induced by stress. Additionally, this mechanism may be targeted to modify behavioral responses to stress.

**Figure 5.**
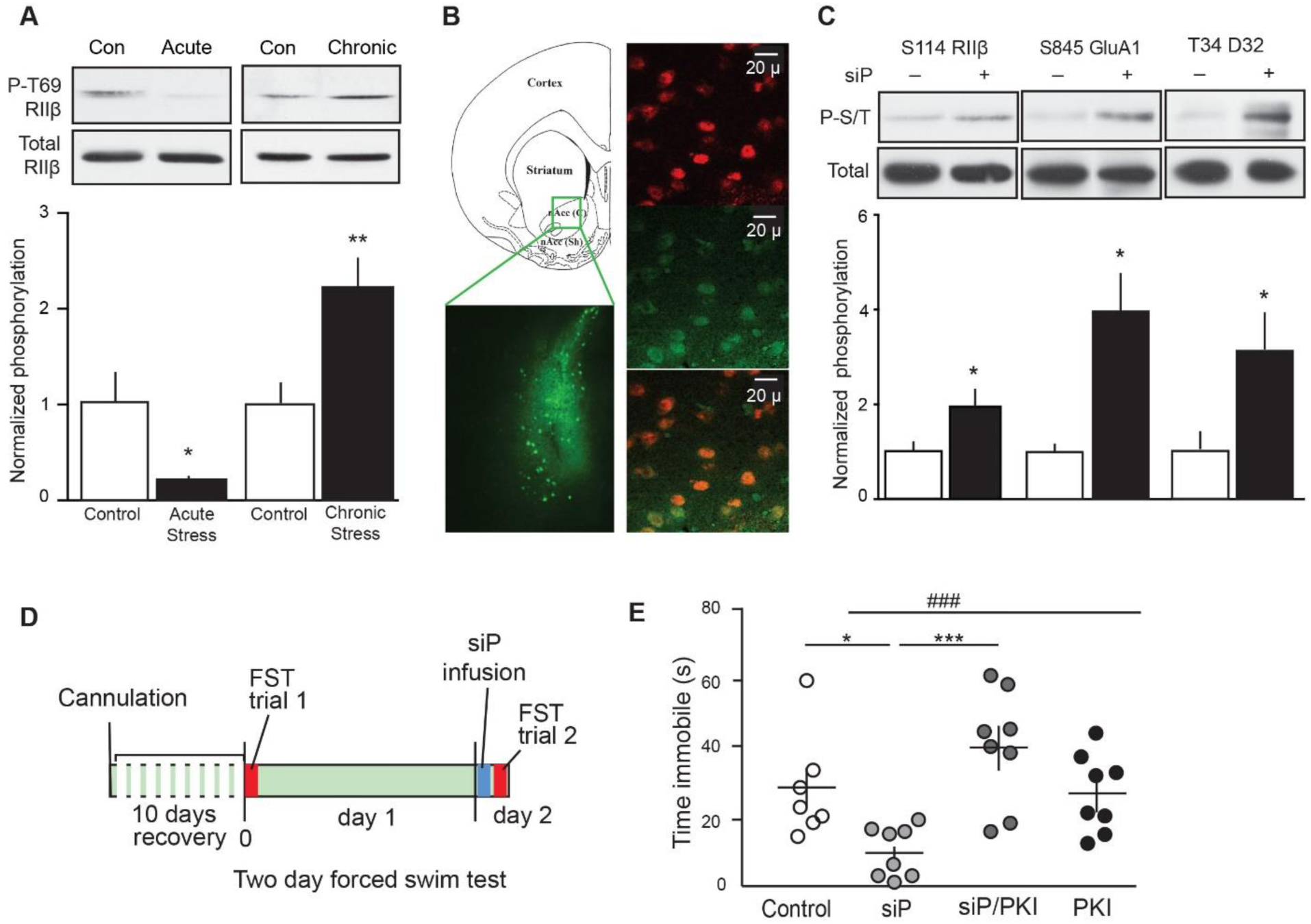
Intra-accumbens targeting of RIIβ regulation improves behavioral responses to stress. (A) Immunoblots of NAc lysate from animals exposed to acute vs. chronic stress (\**p*<0.05, \*\**p*<0.01, unpaired *t*-test, n=4-8). (B) Immunostain of FITC-labeled Thr69 RIIβ siP in nucleus accumbens core. Anatomical diagram shows region of injection and siP detection (left) with labeling of MSNs with nuclear NeuN (red) counterstain and siP (green) in soma and neuropil (right). (C) Immunoblots of lysates from ventral striatum taken 1 h after scrambled (-) vs. RIIβ-siP (+) infusion (\**p*<0.05, unpaired *t*-test, n=6). (D) Depiction of experimental plan for peptide infusion and FST. (E) FST analysis of effects of bilateral intra-accumbens infusions of control (scrambled peptide), RIIβ siP, and/or PKI (^###^*p*<0.001, one-way ANOVA; \**p*<0.05 siP vs. scramble and siP vs. PKI, \*\*\**p*<0.001 siP vs. siP/PKI, Bonferroni multiple comparisons post-hoc analysis, n=7-8). See also Figure S5.

To examine this possibility, bilateral intra-accumbens infusions of the RIIβ siP via cannula were conducted. Delivery of FITC tagged RIIβ siP to ventral striatal core MSNs was confirmed by immunostains (**Figure 5B**). RIIβ siP infusion induced increased PKA-dependent phosphorylation of Ser114 RIIβ, Ser845 GluA1, and Thr34 DARPP-32 *in vivo* (**Figure 5C**), similar to its effects in slices. Incorporating this approach with behavioral studies, animals were subjected to the two-day forced swim test (FST. **Figure 5D**). On day one, animals that had undergone intra-accumbal cannulation were made to swim for 8 min. The following day, subjects were infused with either the RIIβ siP or a scrambled control peptide, then re-exposed to the stress of forced swimming. RIIβ siP-infused subjects spent significantly less time immobile (**Figure 5E, left**). The RIIβ siP effect was completely blocked by co-administration of the PKA inhibitor, PKI. No changes in immobility time were detected when PKI alone was infused. Moreover, none of these infusions had a significant effect on latency to immobility in the FST or locomotor activity (**Figures S5A and S5B**). Immobility was significantly less for animals that received the RIIβ siP during the 4-6 and 6-8 min periods of forced swimming compared to controls receiving scramble peptide (**Figure S5C**). Also, the RIIβ siP’s effects closely mimicked those of the rapid-acting antidepressant, ketamine, administered at sub-anesthetic levels via bilateral intra-accumbal infusion. These findings demonstrate that RIIβ phosphorylation in the NAc mediates behavioral reactions to a stressful environment, and that these responses may be altered by pharmacological targeting of this integrative mechanism.

## DISCUSSION

Emotional salience is imparted upon environmental experiences through integration of cortical glutamatergic and midbrain dopaminergic neurotransmission. The striatum integrates these inputs to mediate reward, motivation, and behavioral flexibility, allowing for learned behavioral responses to rewarding or aversive stimuli (Hurtubise and Howland, 2017). The predominant second messengers that mediate dopaminergic and excitatory glutamatergic activity are cAMP and Ca^2+^, respectively. These signaling modes can act antagonistically by activating protein phosphorylation or dephosphorylation of the same substrate site. However, they may function most critically when working synergistically to invoke downstream effectors that alter neuronal excitability (Vanhoose et al., 2006). For example, integration may occur upstream of PKA, at the level of cAMP metabolism, as both Ca^2+^/CaM-dependent adenylyl cyclase (Wong et al., 1999, Dunn et al., 2009) and phosphodiesterases (Krishnamurthy et al., 2014) can control cAMP/PKA signaling. cAMP/PKA and Ca^2+^ cascades can also converge at key effectors downstream of PKA such as DARPP-32 (Scheggi et al., 2018) and CREB (Steven et al., 2020). Here, we reveal convergence directly upon the RIIβ/PKA_cat_ holoenzyme complex as a fundamental mechanism of glutamate and dopamine integration.

The novel mechanism by which phosphorylation of the RIIβ regulatory subunit governs PKA holoenzyme activation involves the ability of Cdk5-dependent phosphorylation of Thr69 to reduce PKA autophosphorylation of Ser114 RIIβ. Phosphorylation at Ser114 has the net effect of increasing PKA activity by reducing the ability of RIIβ to inhibit PKA_cat_ once activated. Both Thr69 and Ser114 are located within the nonstructural linker region of RIIβ that occurs between the AKAP binding/dimerization domain and cAMP-binding portion of RIIβ. Despite the high level of flexibility in this region, the linker has been demonstrated to play an important role in the overall conformation of the PKA holoenzyme. Holoenzymes containing RIIβ are the most structurally compact compared to those of other regulatory isoforms despite possessing the longest linker sequence of any R subunit (Lu et al., 2020). This feature is conferred by the unique sequence of the RIIβ linker region itself, as substitution of the RIIβ linker into RIIα renders holoenzyme structural density similar to that of PKA_cat_/RIIβ (Vigil et al., 2004). Phosphorylation of the proline-directed Thr69 site may also impart constraints or conformational changes in RIIβ secondary structure that prevent S114 from optimally occupying the catalytic pocket of PKA. Ongoing studies are aimed at solving the structural basis for these effects and better understanding how Ser114 phosphorylation attenuates PKA inhibition. Detailed understanding of the interactions between RIIβ, PKA_cat_ and other protein kinases and phosphatases in the context of AKAP signalisomes (Torres-Quesada et al., 2017) which contribute to synaptic plasticity (Sanderson and Dell’Acqua, 2011), also remains an important area of investigation.

D1-dopamine receptor and glutamatergic NMDAR co-activation enhances striatal plasticity (Yagishita et al., 2014) and is required to achieve corticostriatal LTP in dorsolateral striatum (Hernandez et al., 2016). PKA phosphorylation of DARPP-32 (Calabresi et al., 2000, Yger and Girault, 2011) and GluA1 (Oliveira et al., 2012) both contribute to striatal plasticity. Phosphorylation of Thr34 facilitates plasticity by converting DARPP-32 into a potent PP1 inhibitor, thereby potentiating PKA activity. Phosphorylation at Ser845 GluA1 enhances AMPAR insertion into postsynaptic membranes, a necessary step in LTP. Here, modulation of RIIβ phosphorylation either via NMDAR activation or with an RIIβ siP, resulted in PKA-dependent phosphorylation of GluA1, DARPP-32, and likely other downstream effectors. Importantly, selective reduction of phospho-Thr69 RIIβ markedly improved corticostriatal LTP in ventral striatum, consistent with the known functional contributions of these pathways.

Ventral striatal plasticity is thought to be crucial for coordination of behavioral responses to stressful environments through reinforcement learning and the encoding of rewarding vs. aversive stimuli (Volman et al., 2013). Proper integration of fast and slow neurotransmission may be required for accurate assignment of emotional salience to stimuli and appropriate goal-directed behavioral responses, such as avoidance (Cahill et al., 2014). Here, acute stress caused dephosphorylation of Thr69 RIIβ, and intra-accumbens reduction of Thr69 RIIβ increased struggle in response to re-exposure to forced swim stress. Thus, the regulation of RIIβ phosphorylation and its consequent effects on PKA activity may be an important mechanism mediating stress response behaviors.

Chronic exposure to stress caused an increase in the phosphorylation state of Thr69 RIIβ. Acute stress can improve behavioral and mental acuity (Yuen et al., 2011). However chronic and unpredicted stress can result in a range of deleterious effects including cognitive impairment (Yuen et al., 2012) and is a major predisposing factor for mental illnesses such as major depressive disorder (Scarpa et al., 2020). Elevated levels of stress hormones have been linked to suppression of PKA signaling (Keil et al., 2016). Furthermore, dysregulation of the PKA pathway has been implicated in psychiatric disorders including depression (Yuan et al., 2016). Thus, cAMP/PKA signaling presents an important target for drug therapy. Indeed, antidepressant medications often act either directly or indirectly upon G-protein coupled catecholamine receptors that modulate cAMP/PKA signaling. However, therapies that act at the receptor level often cause unwanted side effects. Thus, more direct manipulation of cAMP/PKA signaling is an attractive alternative approach to antidepressant treatment. If RIIβ phosphorylation can be targeted by small compounds to achieve similar effects as those demonstrated here using the RIIβ siP, perhaps improved health care for mood disorders may be achieved.

## EXPERIMENTAL MODEL AND SUBJECT DETAILS

10-14 week-old male Sprague-Dawley rats (Charles River, strain code 400) were used for most animal experiments and maintained on a 12 h light/dark cycle, single- or double-housed (chronic stress paradigm only) in standard cages, with food and water available *ad libitum* unless otherwise specified (*e*.*g*. food restriction in chronic stress paradigm). For single unit patch clamp recordings, Drd2-EGFP mice in an FVB background were used. All manipulations were approved by UT Southwestern Medical Center and University of Alabama at Birmingham Institutional Animal Care and Use Committees and conducted in accordance with the NIH *Guide for the Care and Use of Laboratory Animals*.

## METHOD DETAILS

### Antibodies and reagents

All chemicals were purchased from Sigma-Aldrich unless otherwise specified. The sequence of the penetratin-tagged RII β-siP and penetratin-tagged scramble control peptides were RQIKIWFQNRRMKWKK-AGAAAGGGTPSKGVNFAEE and RQIKIWFQNRRMKWKK-AAGAGSGAVAFKGANGEA or RQIKIWFQNRRMKWKK-AGAAAGGGAPRKGVNFAEE, respectively. The FITC-tagged siP was synthesized with an N-terminal FITC label

### Protein purification, in vitro phosphorylation, and phosphorylation site identification

Rat recombinant RIIβ was derived as a C-terminal 6XHis fusion by Ni-NTA affinity purification using standard pET vector methods (Novagen). RIIβ was phosphorylated *in vitro* by Cdk5 or PKA using standard conditions (Bibb et al., 2001). Cdk5/p25 was from Millipore, PKA was from NEB. ^32^P incorporation was quantitated by PhosphorImager analysis. All kinetic analyses were conducted under empirically defined linear conditions. PKA activity was measured by phosphorylation of recombinant inhibitor-1 (Nguyen et al., 2007) or DARPP-32 (Bibb et al., 1999). For phosphorylation site identification, preparatively phosphorylated RIIβ was subjected to nano-liquid chromatography tandem mass spectrometry (nano-LC/MS/MS) as previously described (Tassin et al., 2015). Digests were redissolved in 0.1% trifluoracetic acid and loaded onto a ZipTip with C_18_ resin (Millipore) for purification. Analysis was conducted using nanoelectrospray-QSTAR Pulsar I quadrupole time-of-flight tandem mass spectrometry (ESI-Qq TOF MS/MS; MDS Proteomics, Odense, Denmark). Structural model of AKAP/RIIβ dimer interaction was based on published data of AKAP (PDB: 2HWN) (Kinderman et al., 2006) and RIIβ (PDB: 1CX4) (Diller et al., 2001).

### Histology

Immunostains were performed as previously described (Nishi et al., 2008). Briefly, coronal, frozen sections (30 µ) were microwaved in antigen retrieval solution (BioGenex) at 95°C for 10 min and tissue was co-stained for AKAP150 (C-20, Santa Cruz Biotechnology Cat# sc-6445 RRID:AB_2225905; 1:200) and RII β (BD Biosciences Cat# 610626, RRID: AB_397958; 1:2,000). For phospho-T69 RIIβ staining, transverse rat brain cryosections (30 μ) were made. The 1° antibody was used at 1:100 dilution (A phospho-T69 RII β antibody was generated as previously described (Czernik et al., 1997) using the RII peptide GGT*PSKGC (asterisk denotes phospho-Thr); 1:500 2°). For FITC-peptide verification, a cannulated rat was infused with 200 µM FITC-penetratin siP (1 µl over 10 min). After 1 h, brains were collected, placed into a matrix, cut adjacent to the cannula tracts, fixed in O.C.T., and snap-frozen. Cryosections (10 µ) were then stained for FITC (Abcam Cat# ab19224 RRID:AB_732395; 1:400) and counterstained with NeuN (Millipore Cat# ABN78 RRID:AB_10807945; 1:1000).

For staining of primary cultured neurons, embryonic striatal neurons (E18) from Long Evans rats (Charles River Labs) were cultured 14 to 21 days in vitro on 12-mm coverslips, fixed, immunostained for Cdk5 and RII β using the above indicated antibodies and imaged as previously described (Kansy et al., 2004).

### Acute slice pharmacology and quantitative immunoblotting

Slice pharmacology was conducted as previously described (Nishi et al., 1997). Briefly, striatum was microdissected from 350 µ coronal rat brain slices, equilibrated for 1 h, 30°C in Kreb’s buffer with 10 μg/ml adenosine deaminase (Roche). Pharmacological treatments included CP681301 (provided by Pfizer): 0-50 µM, 1h; NMDA: 25 or 50 μM, 5 min; dopamine: 10 µM, 15 min; siP: 10 µM, 1 h; okadaic acid (Cell Signaling): 0.2 or 1 µM, 1 h; and cyclosporin A: 10 µM, 1 h. The NMDA/dopamine co-treatments (NMDA 25 µM, 5 min; DA 10 µM, 15 min) were designed to mimic conditions used to induce striatal LTP (Hernandez et al., 2016). Briefly, conditions included low Mg^2+^ (0.5 mM), and a 30 min tissue recovery in drug-free buffer prior to harvest. Tissue was snap-frozen on dry ice. Immunoblotting was as previously described (Sahin et al., 2006) and analyzed using Image J software (NIH). Data from immunoblots with phosphorylation-specific antibodies were normalized to total protein. Antibodies used included those for PKA regulatory subunits (BD Biosciences Cat# 610626, RRID: AB_397958; BD Biosciences Cat# 610609 RRID:AB_397943; BD Biosciences Cat# 612242 RRID:AB_399565), phospho-S114 RIIβ (BD Biosciences Cat# 612550 RRID:AB_399845), and NMDAR1 (BD Biosciences Cat# 556308 RRID:AB_396353), phospho-S845 GluA1 (PhosphoSolutions Cat# p1160-845 RRID:AB_2492128) and phospho-T34 DARPP-32 (PhosphoSolutions Cat# p1025-34 RRID:AB_2492068), and total GluA1 (Millipore Cat# 05-855R RRID:AB_1587070). Total DARPP-32 antibody was from H. Hemmings (Weill Cornell Medical College). A phospho-T69 RIIβ antibody was generated as previously described (Czernik et al., 1997) using the RII peptide GGT*PSKGC (asterisk denotes phospho-Thr).

### Electrophysiology

Adult rats were anaesthetized by isoflurane (Piramal Healthcare), and brains were placed in ice-cold sucrose saline solution (4°C) containing (in mM): 87 NaCl, 75 Sucrose, 2.5 KCl, 1.25 NaH_2_PO_4_, 7 MgCl_2_, 0.5 CaCl_2_, 25 NaHCO_3_, and 10 glucose (pH 7.4; saturated with 95% CO_2_, 5% O_2_). Parasagittal vibratome slices (350 µ thick) containing prefrontal cortex and NAc core were transferred into saline solution containing 125 NaCl, 2.5 KCl, 1.25 NaH_2_PO_4_, 1.3 MgCl_2_, 2 CaCl_2_, 25 NaHCO_3_, and 25 glucose (pH 7.4; saturated with 95% CO_2_; 5% O_2_) and maintained for 20 min at 30°C, then transferred to room temp. (22-25°C) and allowed 1 h recovery. Slices were transferred to perfusion chambers placed within an upright microscope stage and visualized by infrared differential interference microscopy and a CCD Super Low Luminance camera (KT&C, Co., Ltd). Slices were perfused continuously with oxygenated saline solution (2-3 ml/min).

Extracellular recordings: Field excitatory postsynaptic potential (fEPSP) were obtained at 25°C in the presence of the GABA_A_ antagonist (SR95531, Gabazine, 2 µM) and were evoked by square current pulses (0.2 ms) at 0.033 Hz with a bipolar stimulation electrode (FHC, Bowdoinham, ME) placed at the border of corpus callosum separated by ∼300-500 μ from the recording electrode. The perfusion saline solution was partially modified (in mM): 0.5 MgCl_2_, 1 CaCl_2_, 3.5 KCl. Results were obtained using a stimulus intensity to induce 60% of the maximal fEPSP amplitude taken from the I/O curve of each slice. The paired pulse ratio (PPR) paradigm was applied using different inter-stimulus intervals and PPR changes were measured as PPR = second fEPSP amplitude/first fEPSP amplitude. After recording stable baseline for at least 10 min, LTP was induced by high frequency stimulation (HFS, 4 trains, 100 Hz, 1 s duration, separated by 20 s). fEPSP amplitude was monitored for at least 50 min after HFS to evaluate LTP. All recordings were performed using a Multiclamp 700B amplifier and filtered at 4 kHz and digitized with a Digidata 1440 with pClamp 10 software for data acquisition (Axon, Molecular Devices, LLC, Sunnyvale, CA, USA). Recording pipette was filled with the same extracellular solution as the perfusion bath (2-4 MΩ resistance).

For whole-cell recordings, Drd2-EGFP FVB mice were deeply anesthetized and transcardially perfused with ice-cold cutting artificial cerebrospinal fluid (aCSF) containing (in mM): 87 NaCl, 2.5 KCl, 0.5 CaCl_2_, 7 MgCl_2_, 1.25 NaH_2_PO_4_, 25 NaHCO_3_, 25 glucose, and 75 sucrose, bubbled with 95% O_2_/5% CO_2_. Parasagittal slices were cut transversely at 300 µm using a vibrating blade microtome (VT1200S, Leica Microsystems). Slices were transferred to normal saline solution containing (in mM): 119 NaCl, 2.5 KCl, 2.5 CaCl_2_, 1.3 MgCl_2_, 1.3 NaH_2_PO4, 26 NaHCO_3_, and 20 glucose, at 32°C for 30 min and then allowed to recover for 1 h at room temperature before recordings. Individual slices were transferred to a submerged chamber mounted on a fixed-stage upright microscope (Zeiss Axioskop FS) and continuously perfused at 30°C with normal saline solution containing (in µM): 0 MgCl_2_, 50 picrotoxin, and 10 glycine. GFP-negative D1 cells were visualized by infrared differential interference contrast microscopy with a water-immersion 63X objective (0.9 NA, Zeiss). Current clamp recordings were performed with unpolished pipettes containing (in mM): 140 K-gluconate, 5 KCl, 2 MgCl_2_, 10 HEPES, 4 Mg-ATP, 0.3 Na-GTP, 10 Na_2_-creatine phosphate, 0.2 Na-EGTA, 290-300 mOsm, pH 7.3 (final resistance, 3-4MΩ). LTP of EPSPs was induced by HFS in control and RIIβ siP-treated D1 cells (at -70mV). Slices were incubated in the presence of 10µM control or RIIβ siP for 1 h and washed out prior to recordings. Following LTP, slices were perfused with the NMDAR antagonist D-AP5 (20 μM) and the AMPAR antagonist NBQX (10 μM). All recordings were acquired with Axopatch-200B amplifiers (Molecular Devices), filtered at 2kHz, and digitized at 10 kHz with ITC-18 A/D-D/A interfaces (Instrutech) controlled by custom-written software in G5 PowerMac computers (TI-WorkBench, provided by Dr. Takafumi Inoue).

### Stereotaxic surgeries

Rats were anesthetized with isoflurane and placed in a stereotaxic frame (Kopf Instruments). Bilateral cannulas (3 mm center-to-center distance, cut 8 mm below pedestal, Plastics One) were placed in NAc core (anterior-posterior to bregma +1.5 mm, medial/lateral ±1.5 mm, depth -6.8 mm). Peptides were bilaterally infused 1 week later in awake and mobile animals. Injection volume was 1 µl of peptide (100 µM in PBS, 10% DMSO, per cerebral hemisphere). Animals were allowed to recover for 1 h prior to behavioral testing. Accuracy was confirmed by infusion of methylene blue dye or direct visualization of cannula tracts post-behavioral testing.

### Behavioral Analysis

The Porsolt forced swim test (FST) was conducted as described previously (Castagne et al., 2011). Briefly, one-trial forced swim was used for acute stress, Non-surgerized rats underwent an 8 min swim in an 8-gallon transparent water filled cylinder at 24-25°C. Animals recovered in their home cages for 1 h prior to brain dissection. The NAc was then microdissected for immunoblotting. For two-trial FST, rats recovered from surgical cannulation procedures for 7 days prior to behavioral testing. On day 1, animals were placed in the testing room for 1 h to acclimate. They then underwent an 8 min pre-swim. Video recording was performed from the swimming tank side to observe front and hind limb motion. Awake and mobile animals were bilaterally infused 24 h later with scrambled peptide or siP and/or PKI (1 µl,100 µM), or ketamine (2 µg in 1 µl) at a rate of 0.1 µl/min. They were then placed in a locomotor chamber and locomotor activity was monitored for 30 min in circular donut-shaped locomotor chambers in the dark with a computer-monitored infrared photobeam system (MED-PC, Med Associates). Locomotor counts were defined as sequential adjacent beam breaks. Locomotor testing was followed by a 30 min recovery in home cages, and then 8 min test swims. The last 6 min of the test were scored for latency to immobility and time immobile, with immobility defined as the time animals spent completely motionless.

The chronic unpredictable stress (CUS) paradigm was performed as adapted from Willner and colleagues (Willner, 1997). Animals were pair-housed for 14 days prior to testing. Four pairs of control animals received only regular cage cleaning and handling, while the four pairs of test animals were exposed to 2 stressors per day, chosen pseudorandomly, for 14 days. Stressors included: Cage crowding (4/cage, 2 h), overnight isolation (1/cage, 12 h), overnight food deprivation (12 h), overnight water deprivation (12 h), cage placed at 4°C (45 min), cage placed on orbital shaker (1 h), forced swimming (4 min), reverse lighting for dark cycle (12 h), lights off mid-cycle (3 h), tilted cage (45°, 12 h), and aversive odor exposure (cat urine, 12 h). Animals did not experience any stressor more than 4 times during the testing period and at least 3 days were allowed between any one specific stressor. Brains were harvested 12 h after final stressor.

## QUANTITATION AND STATISTICAL ANALYSIS

### Statistical analysis

Data are reported as mean or normalized mean ± SEM. Statistical analysis was performed using the Student’s *t-*test or Wilcoxon signed-rank test, or one-way analysis of variance (ANOVA) with Bonferroni *post hoc* comparison using GraphPad Prism 6.0 unless stated otherwise. For all experiments, \**p*<0.05, \*\**p*<0.01, \*\*\**p*<0.001 were considered significant. No statistical methods were used to predetermine sample sizes, but sample sizes are similar to those generally employed in comparable studies. Statistical analysis was conducted with n ≥ 4 except for *in vitro* biochemistry (n=3).

## AUTHOR CONTRIBUTIONS

R.T., A.H, D.R.B., W.L. and C.T. conducted experiments, derived and analyzed data. R.T., F.P., A.C., L. P.-M., S.S.T., and J.A.B. contributed to study design, supervision, data interpretation, and manuscript editing. R.T. and J.A.B. wrote the manuscript.

## ACKNOWLEDGEMENTS

We thank H. Ball (UT Southwestern Protein Chemistry Core) for peptides; I. Bowen and the UTSW morphology core for help with microscopy; D. Guzman and S. Birnbaum for technical advice; H. Shu for mass spectrometry, A. Kornev for help with modeling, M. Grey for transgenic mice, and UTSW ARC for help with antibody generation. R.T. is the recipient of the P.E.O. Scholar Award and received support from training grants DA7290 and MH076690. This work was also supported by National Institutes of Health grants to W.L. (NS108508, NS097913), L. P.-M. (NS103089), S.S.T (GM34921) and J.A.B. (MH083711, DA033485, MH116896, MH126948).

