## Supplemental Figures 1-5 for "Integrated Regulation of PKA by Fast and Slow Neurotransmission in the Nucleus Accumbens Controls Plasticity and Stress Responses"

### INVENTORY OF SUPPLEMENTAL INFORMATION

**Supplemental Figures:**

**Figure S1** Related to data in Figure 1 and complementing Figure 2

**Figure S2** Related to data in Figure 3

**Figure S3** Data complementing Figure 4

**Figure S4** Data complementing Figure 4

**Figure S5** Data complementing Figure 5

### SUPPLEMENTAL FIGURES

Figure S1, Thomas, et al.

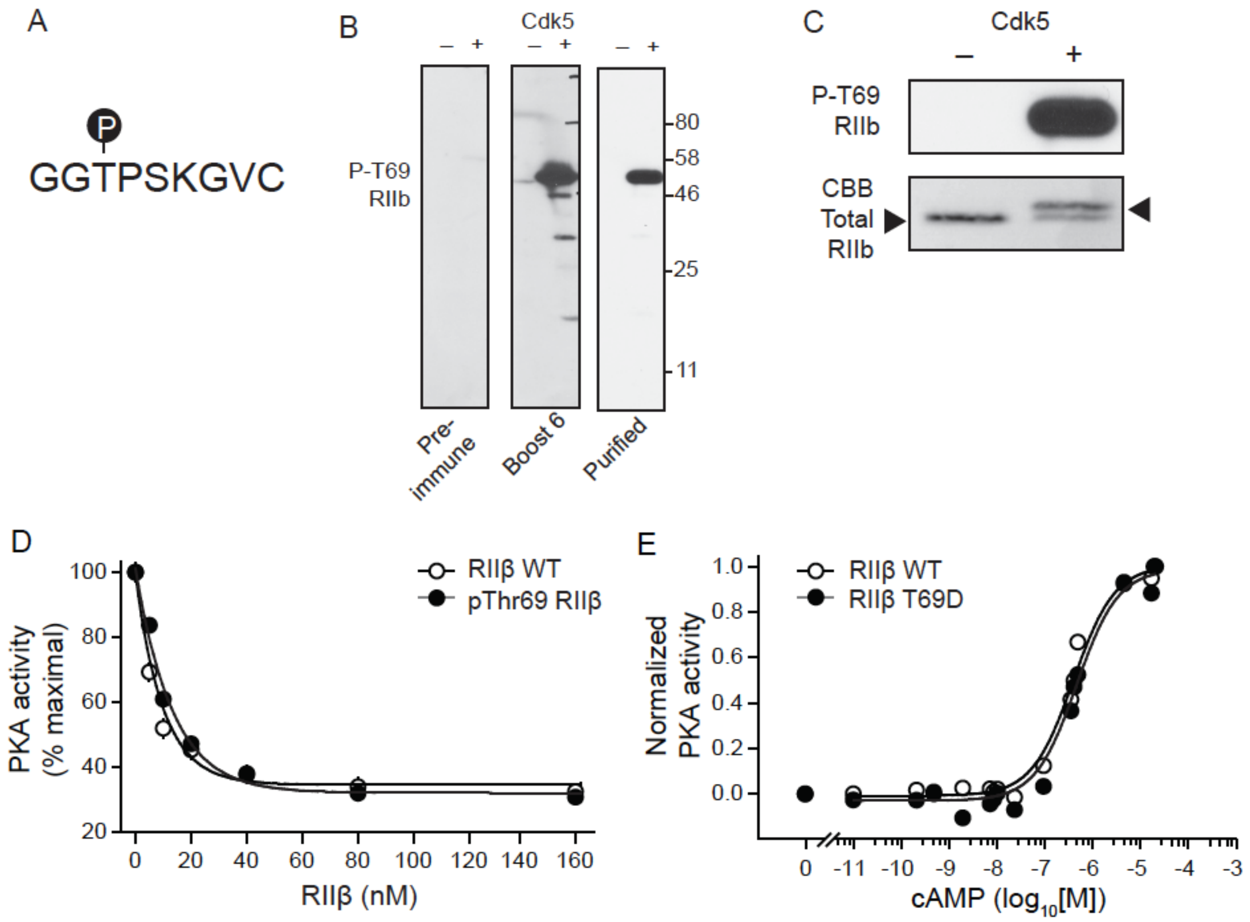


**Figure S1. Verification of phosphorylation-state specificity of phospho-T69 RIIβ antibody and assessment of the effect of phopsho-Thr69 on RIIβ/PKAcat or RIIβ/cAMP interactions, related to Figures 1 and 2.**
(A) Amino acid sequence of the phospho-peptide used to generate the polyclonal anti-phospho-T69 RIIβ antibody. (B) Immunoblots of unphosphorylated (-) and *in vitro* phosphorylated (+) recombinant RIIβ with serum samples (preimmune, left; post-antigen boost 6, middle) and affinity purified (right) phospho-T69 RIIβ antibody. (C) Immunoblot (top) of unphosphorylated (-) and *in vitro* phosphorylated (+) recombinant RIIβ using phospho-T69 RIIβ antibody compared to total pure Coomassie Blue stained protein (bottom). Phosphorylation of RIIβ caused an upward shift in SDS-PAGE mobility. (D) Plot of PKA inhibition by dephospho- vs. phospho-T69 RIIβ. (E) Analysis of cAMP-dependent activation of PKA complexed with WT vs. T69D RIIβ. Data represent means ± S.E.M., n=4.

Figure S2, Thomas, et al.

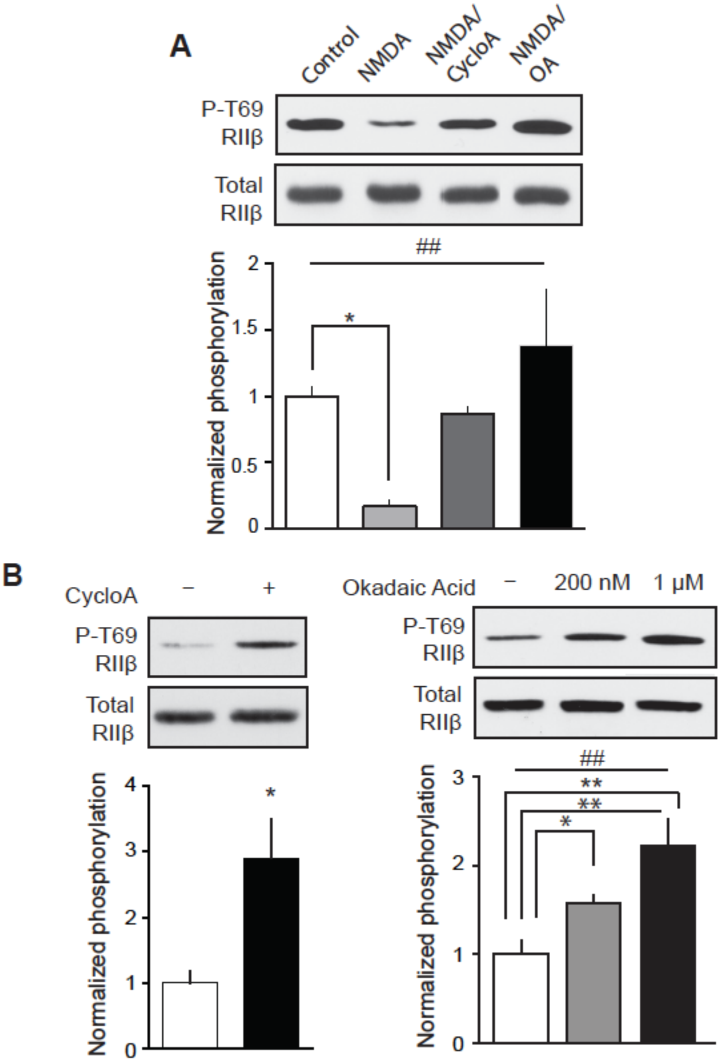


**Figure S2. Regulation of phospho-Thr69 RIIβ by NMDA is mediated through protein phosphatases PP2A, 2B, or 1, related to Figure 3.** (A) Quantitative immunoblot analysis of lysates from striatal slices treated with NMDA (25 μM, 5 min), in the absence or presence of the indicated protein phosphatase inhibitors cyclosporin A (CycloA, 10 μM, 1 h) or okadaic acid (OA, 1 μM, 1 h: ##$p$<0.005, one-way ANOVA; *$p$<0.05, multiple comparisons, control vs. NMDA, n=4-6). (B) Effects of protein phosphatase inhibition on basal T69 RIIβ phosphorylation state (CycloA, 10 μM, 1 h: *$p$<0.05, Student's unpaired $t$-test, n=4-5) and (OA, 200 nM and 1 μM, 1h: ##$p$<0.005, one-way ANOVA; **$p$<0.005, multiple comparisons, control vs. 1 μM OA, n=4-5). All data are normalized means ± S.E.M.

Figure S3, Thomas, *et al.*

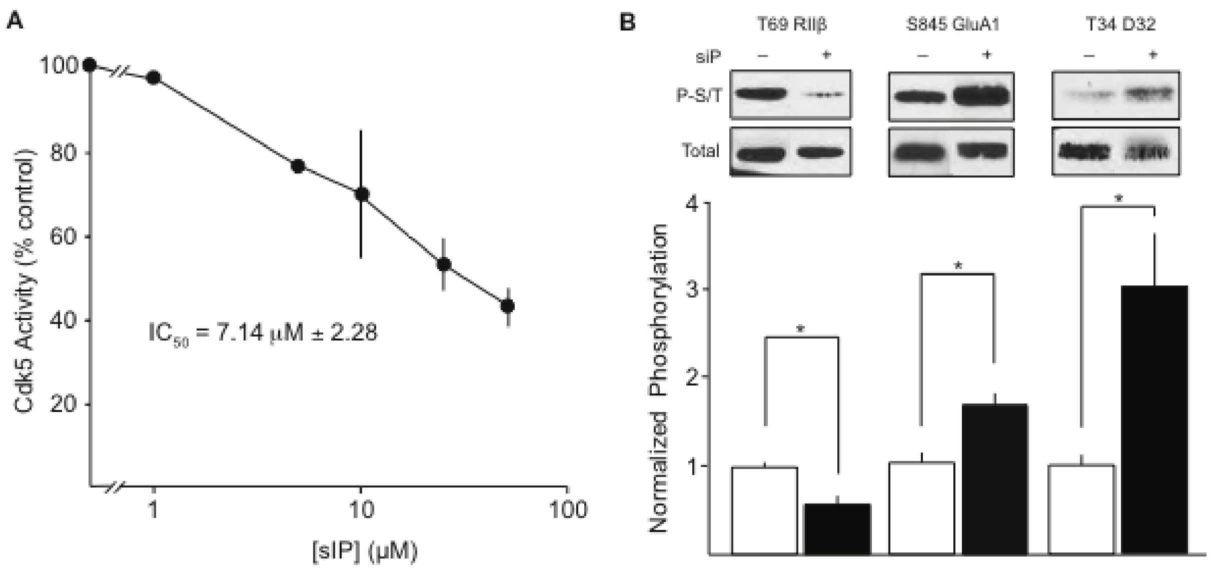


**Figure S3. Inhibition of Thr69 RIIβ phosphorylation by siP *in vitro* and in intact brain, related to Figure 4.**

(A) Inhibition curve for phosphorylation of RIIβ by Cdk5 *in vitro*. Data represent means ± S.E.M. (B) Immunoblot of analysis NAc slice lysates for effect of RIIβ siP treatment (10 μM, 1 h) in the presence of dopamine (10 μM, 15 min) on phospho- (P-S/T) Thr69 RIIβ, Ser845 GluA1, and Thr34 DARPP-32 (*$p$<0.05, unpaired t-test, n=6). Data represent normalized means ± S.E.M.

Figure S4, Thomas, *et al.*

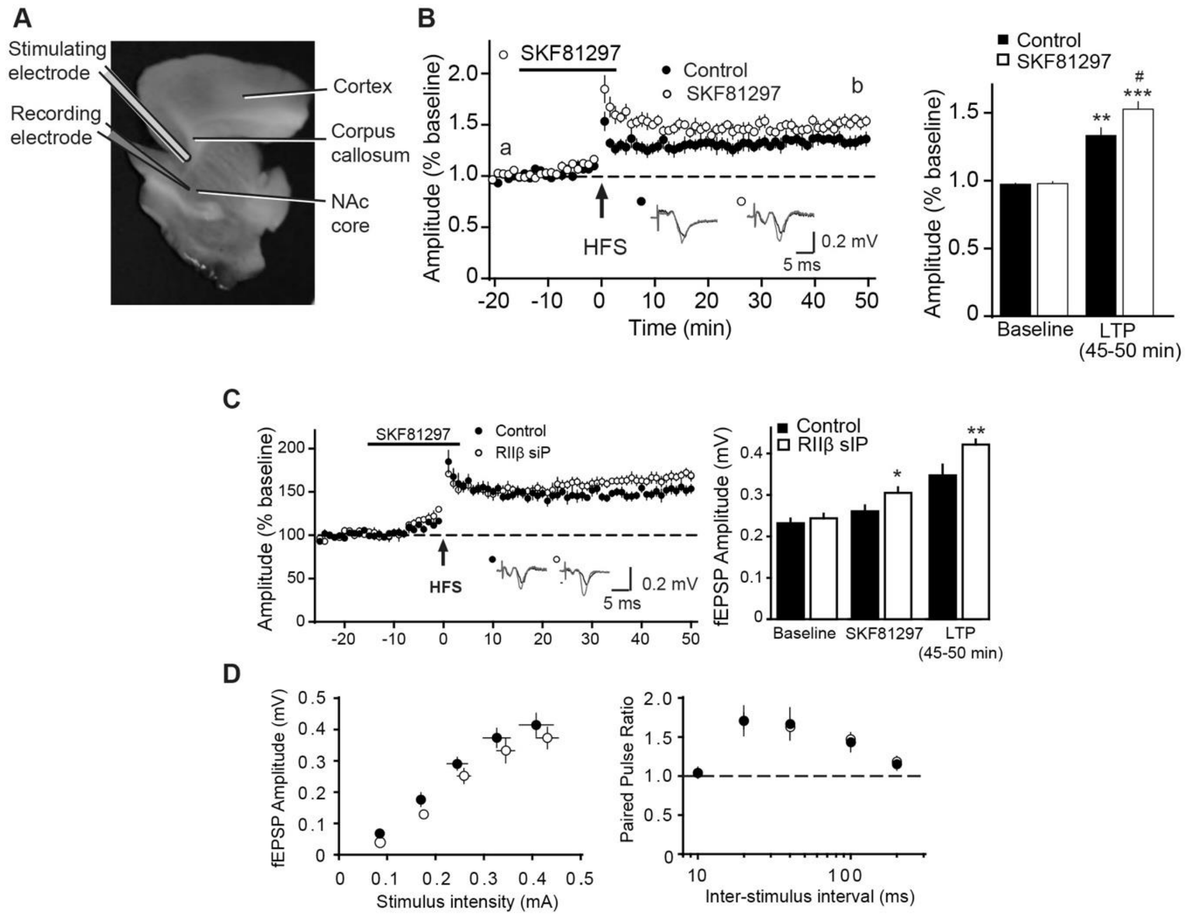


**Figure S4. Positioning of electrodes for field recordings, I/O curves, and PPR, related to Figure 4.**

(A) Depiction of electrode placement for ventral striatal plasticity studies. (B) Effects of the D1 agonist SKF81297 (2 µM) on NAc LTP. Plot of fEPSP amplitudes with sample traces from a vs. b are shown (left) with summary plot (right), $^{**}p<0.01$, $^{***}p<0.001$, basal vs. LTP; $\#p<0.05$ control vs. SKF, unpaired *t*-test, n=7. (C) RIIβ siP effects on HFS-induced LTP (left) in the presence of D1 agonist, SKF81297 with summary plot (right), $^{*}p<0.05$, basal vs. SKF; $^{**}p<0.01$, basal vs. RIIβ siP LTP, unpaired *t*-test, n=8. (D) I/O curves and PPR for slices undergoing siP/SKF81297 protocol. These data correlate with data shown in panel C. Data represent means ± S.E.M.

Figure S5, Thomas, *et al.*

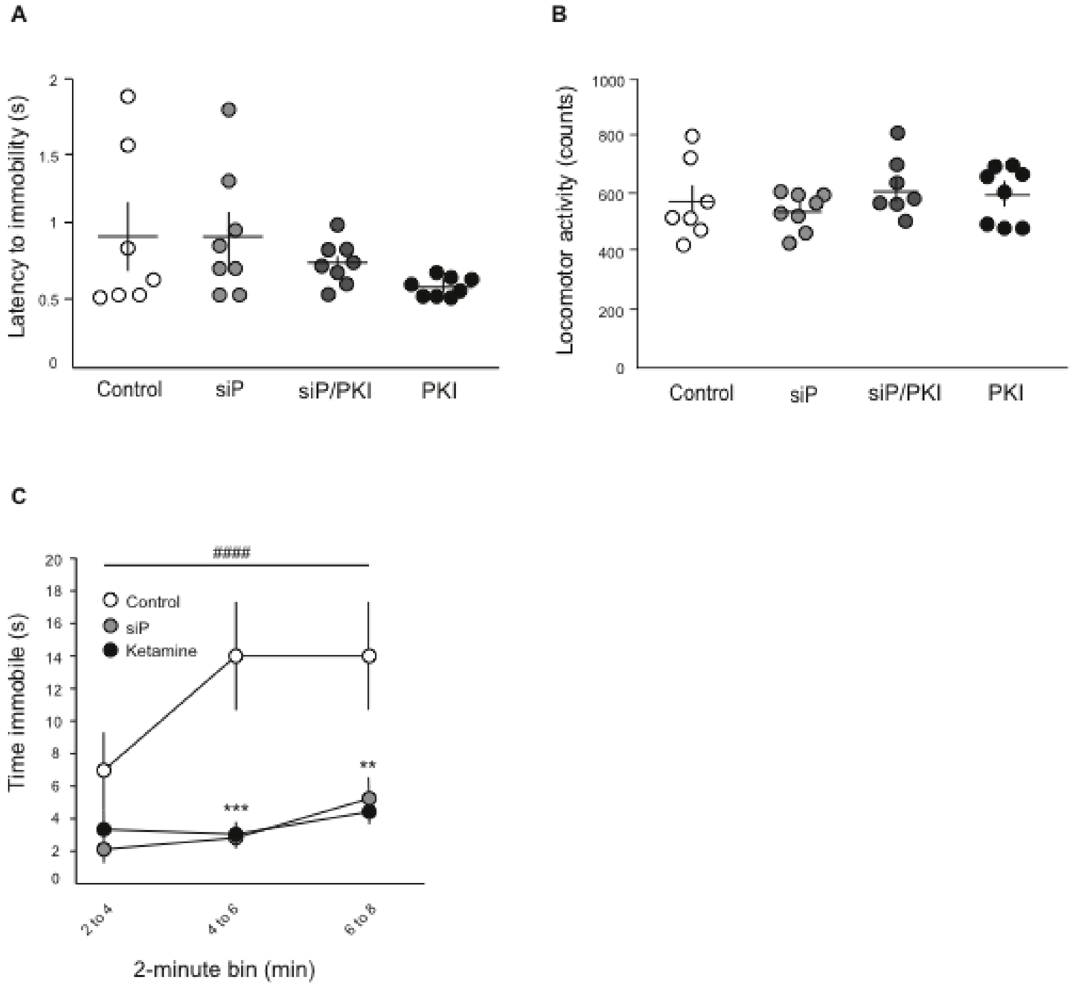


**Figure S5. Stress response behavior in animals following peptide infusion, related to Figure 5**

(A) Analysis of latency to immobility and (B) locomotor effects of bilateral intra-accumbens peptide infusion of control (scrambled), RIIβ siP, and/or PKI. Motor activity was recorded over the first 30 min following infusion. (C) Immobility time binned (2-minute), comparing animals subjected to bilateral intra-accumbens infusion with scramble control peptide, siP (1 µl,100 µM), or ketamine (2 µg in 1 µl). Bins 4 to 6, and 6 to 8 were significant (####$p$<0.0001, two-way ANOVA; ***$p$<0.001 for bin 4 to 6, scramble peptide vs. RIIβ siP and scramble vs. ketamine, **$p$<0.005 for bin 6 to 8, scramble vs. RIIβ siP and scramble vs. ketamine, multiple comparisons, n=6-8). Data represent means ± S.E.M.
